# Biogenic flavonoid capping converts a cytotoxic Carica papaya fraction into a selective, cross-serotype dengue entry inhibitor

**DOI:** 10.64898/2026.09.26.754612

**Authors:** D. Madan Kumar, R. Raj Bharath, Gowtham Palanisamy, G. Devanand Venkatasubbu, Tanay Bhatt

**Author notes:** Corresponding authors. R. Raj Bharath,; Tanay Bhatt.

## Abstract

Plant-derived flavonoids show measurable anti-dengue activity in cell culture, but their translational value is limited by a narrow therapeutic window: the concentrations that inhibit the virus approach or exceed those that are cytotoxic. Whether nanoparticle formulation can resolve this constraint, rather than simply add potency, remains untested for a chemically defined fraction. Here we show that biogenic silver nanoparticle (AgNP) formation using a flavonoid-enriched fraction of Carica papaya inverts an unusable selectivity profile into a viable one. The unformulated fraction was cytotoxic below the concentrations required for antiviral activity: its 50% cytotoxic concentration (CC₅₀ = 134.9 µg/mL) lay below its 50% effective concentration against dengue virus serotype 2 (DENV-2; EC₅₀ = 254.4 µg/mL), giving a Selectivity Index (SI) of 0.53. Using the same flavonoids as sole reducing and capping agents produced AgNPs (Z-average 128.4 nm; PDI 0.232; zeta potential −28.4 mV) that moved both parameters simultaneously. CC₅₀ rose approximately 8.6-fold to 1,165.74 µg/mL while EC₅₀ fell approximately 8.8-fold to 29.05 µg/mL, raising the SI to 40.12, an approximately 75-fold shift. Time-of-addition analysis localised the effect to the extracellular phase: inhibition was significant under pre-treatment and co-treatment but not after viral adsorption, identifying the AgNPs as entry inhibitors rather than replication inhibitors. Consistent with a serotype-independent physical mechanism, AgNP treatment at 30 µg/mL reduced viral RNA across all four serotypes, using inocula standardised against WHO-traceable NAAT reference reagents. These findings identify capping chemistry, rather than silver content alone, as a determinant of the therapeutic window in phytosynthesised nanoantivirals.

## INTRODUCTION

Dengue fever is among the most rapidly spreading mosquito-borne viral diseases globally, with an estimated 390 million infections occurring annually across more than 129 endemic countries, of which approximately 96 million manifest clinically [1]. The clinical spectrum ranges from a self-limiting febrile illness to life-threatening complications including plasma leakage, severe haemorrhage, multi-organ impairment and death [2,3]. Secondary heterologous infection is strongly associated with antibody-dependent enhancement (ADE), which amplifies disease severity and complicates both therapeutic and prophylactic strategies [4]. Despite decades of research there is still no specific antiviral therapy for dengue, and treatment remains supportive [5]. The first licensed vaccine, CYD-TDV (Dengvaxia), represented a milestone, but post-licensure evidence of elevated severe dengue risk in seronegative recipients led the WHO Strategic Advisory Group of Experts to restrict its use to individuals with confirmed prior infection [6]. The therapeutic gap therefore remains open, and it is widest for agents that could act across all four co-circulating serotypes.

Plant-derived flavonoids have been proposed repeatedly as candidates to fill this gap. Specific flavonoids inhibit DENV replication in cell culture, with quercetin showing measurable activity against DENV-2 [7,8], and Carica papaya L. has a long ethnomedicinal record in the management of dengue-associated thrombocytopaenia, with phytochemical studies identifying flavonoids and flavonoid glycosides related to the quercetin and kaempferol families in its leaves [9]. Yet the translational record of these preparations is poor, and the reason is not usually an absence of antiviral activity. Crude polyphenolic mixtures are themselves cytotoxic at therapeutically relevant concentrations, and are further limited by poor aqueous solubility, low bioavailability of free flavonoids and physicochemical instability in biological media [8]. The decisive parameter is therefore not potency in isolation but the separation between the concentration that inhibits the virus and the concentration that harms the host cell, that is, the therapeutic window, expressed in vitro as the Selectivity Index (SI = CC₅₀ / EC₅₀).

This distinction is consequential because the SI is inconsistently reported in the phytochemical and phytosynthesised nanoparticle antiviral literature. Many studies report an effective concentration without the paired cytotoxicity determination performed under the same exposure conditions, leaving it unresolved whether the observed antiviral effect occupies a usable window at all. Where a preparation’s CC₅₀ falls below its EC₅₀, an apparent reduction in viral load may in part reflect loss of viable host cells rather than genuine antiviral selectivity. Establishing whether a formulation strategy can move a preparation from that regime into a selective one is a different question from establishing that it improves potency, and it is the question this study addresses.

Green synthesis of silver nanoparticles (AgNPs) offers a mechanistically plausible route to do so. Silver nanoparticles are recognised for broad-spectrum antimicrobial and antiviral properties, interacting with biological membranes, viral envelope glycoproteins and nucleic acids [10]. The feature that distinguishes biogenic synthesis from conventional chemical routes is that the plant-derived phenolics and flavonoids act simultaneously as reducing agents, converting Ag⁺ to Ag⁰, and as stabilising capping agents that prevent agglomeration while retaining their own biological activity [11]. The capping layer is therefore not inert packaging but a functional surface. This raises a possibility that has not been tested directly, namely that formulation acts on both terms of the selectivity ratio at once: multivalent presentation of flavonoids on a nanoparticle scaffold may lower the effective concentration required for antiviral activity, while sequestration of free phytochemicals within the capping corona and moderated silver-ion release may raise the concentration at which host-cell toxicity appears. If so, the resulting improvement in SI would exceed what either effect could produce alone.

Testing this requires a chemically defined starting material. Earlier work showed that AgNPs synthesised from a methanolic crude extract of C. papaya inhibited DENV-2 replication in vitro [12], but crude, unfractionated extracts carry phytochemical heterogeneity that limits reproducibility and prevents attribution of antiviral activity to a defined compound class. We therefore used systematic liquid–liquid partitioning to isolate a flavonoid-enriched chloroform fraction and employed that fraction as the sole reducing and capping agent, so that any change in the selectivity profile can be attributed to the formulation step rather than to compositional drift between preparations.

Accordingly, this study aimed to: (i) isolate and quantify a flavonoid-enriched fraction of C. papaya by systematic liquid–liquid partitioning; (ii) synthesise and characterise biogenic AgNPs using this fraction as the sole reducing and stabilising agent; (iii) determine the acute cytotoxicity (CC₅₀) and antiviral efficacy (EC₅₀) of both the unformulated fraction and the nanoformulation against DENV-2 in Vero cells under matched exposure conditions, and compare their Selectivity Indices; (iv) resolve the stage of the viral life cycle at which the nanoformulation acts, using a time-of-addition design; and (v) screen the nanoformulation for activity across DENV-1 to DENV-4 at a standardised concentration in order to establish whether the mechanism identified is serotype-independent.

## 2. METHODOLOGY

### 2.1 Plant Material Collection and Authentication

Fresh, healthy leaves of Carica papaya L. were collected from department of Horticulture, SRM college of agriculture sciences, SRMIST. To ensure the scientific integrity of the study, the plant specimen was authenticated by a taxonomist at National Institute of Siddha, Ministry of AYUSH under the voucher specimen number NISMB8212025.

### 2.2. Processing and Extract Preparation

The collected leaves were thoroughly washed under running tap water to remove surface contaminants, followed by a final rinse with double-distilled water. The cleaned leaves were shade-dried at room temperature to prevent thermal degradation of the bioactive constituents, then pulverized into a coarse powder using a mechanical grinder. Approximately 40 g of the powdered leaf material (corresponding to a dry weight of 38.71 g, used in the yield calculation below) was loaded into a Soxhlet apparatus, and extraction was performed using 500 mL of 70% ethanol (Figure S1). Extraction continued for 52 hours, until the solvent in the siphoning tube became colourless, ensuring exhaustive extraction of the phytoconstituents [13]. The resulting ethanolic extract was filtered through Whatman No. 1 filter paper, and the filtrate was concentrated using a rotary vacuum evaporator at 40°C - a temperature selected to facilitate solvent removal under reduced pressure while preserving the structural integrity of the thermolabile flavonoids.

The percentage yield for the Soxhlet extraction process was calculated using the following gravimetric equation: Yield (%) = (“Weight of extract obtained” / “Weight of dry sample used”) ×100

### 2.3. Qualitative Phytochemical Screening

The 70% ethanolic crude extract was subjected to preliminary qualitative screening to profile the major secondary metabolite classes. The screening targeted eight phytochemical groups: flavonoids, alkaloids, phenols, saponins, tannins, terpenoids, glycosides, and steroids using standard analytical procedures [14,15].

### 2.4. Liquid-Liquid Partitioning and Fractionation

To isolate bioactive constituents by polarity, the crude extract was subjected to liquid– liquid partitioning using a separating funnel. Briefly, 10 g of crude ethanolic extract was dissolved in 50 mL of hydroalcohol (70% ethanol) to form a mother solution, which was partitioned sequentially with solvents of increasing polarity at a 1:1 (v/v) ratio. At each step, the mixture was shaken vigorously and allowed to stand until distinct layers formed.

The solution was first partitioned with 50 mL of n-hexane to remove non-polar lipids (hexane layer collected; aqueous-alcohol phase retained), then with 50 mL of chloroform to enrich flavonoid aglycones (chloroform layer collected), and finally with 50 mL of n-butanol. All fractions (hexane, chloroform, butanol) and the final aqueous residue were concentrated by rotary vacuum evaporation at 40°C.

### 2.5. Determination of Total Flavonoid Content (TFC)

The quantification of flavonoids in all the fractions was performed using the aluminum chloride (AlCl3) colorimetric method. A 0.5 mL aliquot of all four fractions (1 mg/mL) was mixed with 1.5 mL of ethanol, 0.1 mL of 10% AlCl3, and 0.1 mL of 1 M potassium acetate. The final volume was adjusted to 5 mL with distilled water. After incubation for 30 minutes at room temperature, the absorbance was recorded at 415 nm. Results were extrapolated from a Quercetin standard curve (R² = 0.99) and expressed as mg of Quercetin Equivalents per gram of dry extract (mg QE/g) [16].

### 2.6. Green Synthesis of Silver Nanoparticles (AgNPs)

The flavonoid-enriched chloroform fraction of Carica papaya served as the reducing and stabilizing agent for AgNP synthesis. A 1 mM aqueous solution of Silver Nitrate (AgNO3) was prepared by dissolving 0.169 g of AgNO3 in 1000 mL of deionized water [17]. For synthesis, 10 mL of the C. papaya flavonoid-rich fraction was added dropwise to 90 mL of the 1 mM AgNO3 solution (1:9 ratio), selected to provide an optimal concentration of capping agent and minimize agglomeration. The reaction mixture was stirred continuously at room temperature for 24 hours. Successful AgNP formation was confirmed visually by a colour change from pale yellow to deep reddish-brown [12].

### 2.7. Characterization of Synthesized AgNPs

#### 2.7.1. Morphological and Elemental Analysis (FE-SEM with EDS)

Surface morphology was examined by Field Emission Scanning Electron Microscopy (FE-SEM); samples were prepared by drop-casting the nanoparticle suspension onto a carbon-coated copper grid and drying under vacuum. Energy Dispersive X-ray

Spectroscopy (EDS) was used to determine elemental composition, with the localized silver signal (∼3 keV) confirming metallic silver purity [18].

#### 2.7.2. Particle Size and Surface Charge Analysis (DLS and Zeta Potential)

Hydrodynamic diameter and Polydispersity Index (PDI) were determined by Dynamic Light Scattering (DLS) using a Zeta Particle Size Analyzer. Colloidal stability and surface charge were evaluated by zeta potential, with higher absolute magnitude indicating greater electrostatic repulsion conferred by the flavonoid capping layer [18].

#### 2.7.3. Functional Group Identification (FTIR-ATR)

FTIR spectroscopy was conducted in Attenuated Total Reflectance (ATR) mode to identify functional groups responsible for AgNP reduction and stabilization. Samples were placed directly on the diamond ATR crystal, and scans were recorded across 4000– 400 cm⁻¹. Shifts in absorption bands particularly those of hydroxyl (–OH) and carbonyl (C=O) groups were used to confirm interaction between plant flavonoids and the silver surface [18].

### 2.8. Cell and Viral Culture

All cell culture maintenance, viral propagation, WST-1 cytotoxicity assays, and in vitro neutralization assays were conducted within the biocontainment facilities (BSL-2) at the National Centre for Biological Sciences (NCBS), Bengaluru. C6/36 (mosquito-derived) and Vero (mammalian-derived) cell lines were used for virus propagation and antiviral testing, respectively. C6/36 cells were cultured in Dulbecco’s Modified Eagle Medium (DMEM) supplemented with 10% Fetal Bovine Serum (FBS) and maintained at 28°C, under a 5% CO2 atmosphere. Vero cells were maintained in DMEM with 10% FBS at 37°C under a 5% CO2 atmosphere.

#### 2.8.1. Obtaining DENV serotypes

Characterised dengue virus serotype 1, 2, 3, and 4 were obtained from the NCBS Biorepository facility. Viral titer (U/mL) was quantified using NAAT based secondary reference reagents, traceable to the WHO primary international reference reagent [19]. Working viral stocks were standardized such that untreated infection of Vero cells yielded Ct values of approximately 20–25 at 24 hours post-infection, providing a consistent infectious challenge across assays. Viral stocks were stored at −80°C until use in the antiviral assays.

### 2.9. Cytotoxicity Evaluation (WST-1 Assay)

Cytotoxicity of the flavonoid-rich fraction and AgNPs on Vero cells was evaluated using the WST-1 (Water-Soluble Tetrazolium) colorimetric assay. Vero cells were cultured in 96-well plates at 70-80% confluency, then exposed to fresh medium containing varying concentrations of test compounds (0.6–5000 µg/mL) at a standardized volume of 100 µL per well. Cells were exposed for 24 hours to allow manifestation of cytotoxic effects.

Post-treatment period, 10 µL of WST-1 reagent was added per the manufacturer’s protocol, and plates were incubated for 30 minutes. Absorbance was measured at 450 nm (primary) and 600 nm (reference). Cell viability was calculated as: Cell Viability (%) = [Absorbance of Sample] / [Absorbance of Control] Χ 100.

The 50% cytotoxic concentration (CC50) values were calculated by fitting the concentration vs. response data to a non-linear regression curve using the four-parameter logistic (4PL) model with a variable slope.

### 2.10. In Vitro Time-of-Addition Antiviral Assays

To comprehensively determine the mechanism of action and the specific stage of the viral life cycle targeted by the formulations, a time-of-addition study was conducted encompassing pre-treatment, co-treatment, and post-treatment assays. All experiments were performed in 24-well plates containing ∼ 80% confluent Vero cells, incorporating respective virus-only and cell-only controls.

#### 2.10.1. Pre-Treatment Assay

To evaluate dose-dependent direct virucidal potential, viral aliquots were incubated with varying concentrations (9.375–1200 µg/mL) of the test compounds at 37°C for 1 hour, allowing for direct interaction between the formulations and viral particles. Following incubation, the pre-treated virus–compound mixtures were inoculated onto the Vero cells and incubated at 37°C for 2 hours to facilitate viral adsorption. The inoculum was then aspirated, cells were washed, and fresh maintenance medium was added. The plates were subsequently incubated for 24 hours at 37°C in a 5% CO₂ atmosphere.

#### 2.10.2. Co-Treatment Assay

To assess interference during viral attachment and entry, a single standardized inhibitory concentration of the test compounds (determined based on the pre-treatment assay results) was utilized. The virus and formulations were mixed and immediately added to the Vero cell monolayers without prior incubation. This mixture was left on the cells for 2 hour at 37°C. Subsequently, the inoculum was aspirated, fresh maintenance medium was added, and the plates were incubated for 24 hours at 37°C with 5% CO₂.

#### 2.10.3. Post-Treatment Assay

To investigate the effect of the formulations on post-entry viral replication, Vero cells were first inoculated with the virus alone and incubated for 2 hour at 37° C to allow for viral adsorption. Following this infection period, the viral inoculum was aspirated, and the cells were washed. Fresh maintenance medium containing the standardized inhibitory concentration of the test compounds was then added to the cells, followed by a 24-hour incubation.

### 2.11. Quantitative Real-Time PCR (RT-qPCR) Analysis

To quantify the antiviral effect of each formulation on DENV replication, the viral RNA load was measured utilizing RT-qPCR assay. Initially, total RNA was extracted from the Vero cells collected from the neutralization assay using the column based NeoDx RNA Extraction Kit (Catalogue No: #NDX-RNA-021), as per manufacturer’s instructions. The quantitative amplification was subsequently performed on a Bio-Rad RT-qPCR (Model CFX96) instrument. The reaction mixture was prepared using the Luna Probe One-Step RT-qPCR 4X Mix with UDG (Catalogue No: M3019S). Serotype specific primers and probes for various DENV serotypes utilized for this assay were adapted from a previously validated diagnostic protocol [20] and are detailed in Table S1. Cycle threshold (Ct) values were recorded, and percent viral infection relative to the virus-only control was calculated using the comparative-Ct method: Percent Viral Infection (%) = 2(–ΔCt) × 100, where ΔCt = Ct (treated) – Ct (virus-only control) EC₅₀ values for each formulation were derived by fitting percent viral infection against concentration to a four-parameter sigmoidal (variable-slope) dose-response model in GraphPad Prism software.

### 2.12. Statistical Analysis

All data from the WST-1 and RT-qPCR-based viral infection assays were obtained from three independent replicates and expressed as mean ± SEM. CC₅₀ and EC₅₀ were determined by non-linear regression using a 4-parameter logistic (4PL) curve fit. The Selectivity index (SI) of each formulation was calculated as: Selectivity Index (SI) = CC₅₀ / EC₅₀

Two-way ANOVA, followed by Tukey’s multiple comparisons test, was used to compare experimental groups. All processing and statistics were performed using GraphPad Prism software. A p-value of ≤ 0.05 was considered statistically significant. Significance levels are denoted in figures as *p ≤ 0.05, **p ≤ 0.01, ***p ≤ 0.001, and ’ns’ for non-significant.

## 3. RESULTS

### 3.1 Extraction Yield

From an initial starting dry sample weight of 38.71 g, a total crude extract mass of 12.00 g was recovered following complete rotary solvent evaporation. Utilizing the gravimetric formula, the extraction protocol achieved a final mass yield of 31.00 %.

### 3.2 Qualitative Phytochemical Screening

To profile the classes of secondary metabolites, the crude extract was analysed qualitatively. The screening results, summarised in Table S2, demonstrated a highly selective phytochemical composition dominated by polyphenolic compounds.

### 3.3 Phytochemical Quantification and Total Flavonoid Content (TFC)

To quantify the metabolic profile of the four leaf fractions (aqueous, hexane, chloroform and butanol) of Carica papaya, a quercetin standard calibration curve was constructed, showing linearity at 415 nm with a regression equation of y = 0.0008x + 0.0731, R² = 0.99 (Figure S2). Total flavonoid content differed markedly across the fractions, rising from 13.71 ± 2.69 mg QE/g in the hexane fraction and 23.92 ± 2.03 mg QE/g in the aqueous residue to 34.33 ± 0.44 mg QE/g in the butanol fraction and 55.79 ± 1.94 mg QE/g in the chloroform fraction. The chloroform fraction yielded the highest total flavonoid content and was therefore carried forward both for antiviral evaluation and as the reducing and capping agent for AgNP synthesis.

### 3.4 Synthesis and Characterization of AgNPs

#### 3.4.1 UV-Vis Spectroscopy and Visual Observation

Following addition of the chloroform fraction to AgNO₃, the synthesis of AgNPs was initially confirmed by the visible color transition of the reaction mixture from pale yellow to deep brownish-red. The absorption profile of the precursor flavonoid-rich fraction exhibited a sharp, high-intensity band at λmax = 287 nm (Figure 1A), which is characteristic of the electronic transitions (π → π* and n → π*) associated with the aromatic rings and carbonyl groups of polyphenolic constituents [21]. After synthesis, the successful formation of AgNPs was definitively confirmed by the emergence of a prominent Surface Plasmon Resonance (SPR) peak at λmax = 417 nm (Figure 1B); its narrow, symmetric profile and the absence of a secondary peak beyond 600 nm are consistent with a monodispersed, non-aggregated population [22].

**Figure 1.**
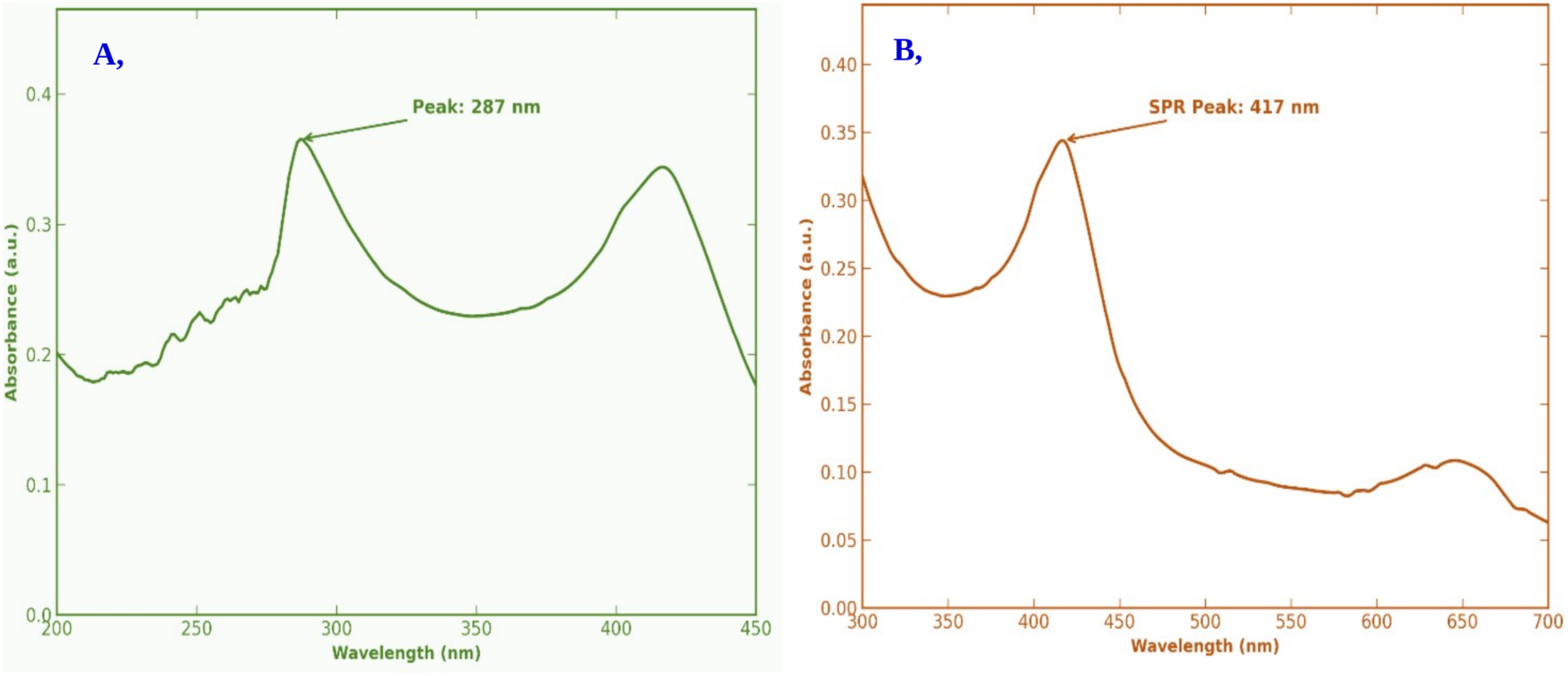
Optical confirmation of AgNP formation. (A) UV-visible absorption spectrum of the flavonoid-enriched chloroform fraction of C. papaya, showing the polyphenol band at λmax = 287 nm. (B) Spectrum after reaction with AgNO₃, showing the surface plasmon resonance band at λmax = 417 nm.

#### 3.4.2 FTIR-ATR Analysis

To identify the functional groups mediating reduction and capping, FTIR-ATR was performed on the extract and the AgNPs (Figure 2). The spectrum of the flavonoid extract is characterized by intense absorption bands at 3365.78 cm-1 (O-H stretching of phenols), 2920.23 cm-1and 2852.72 cm-1 (aliphatic C-H stretching), and 1730.15 cm-1 (C=O stretching of carbonyl groups). After synthesis: the O–H band shifted to 3253.91 cm-1 (bathochromic shift), and the C=O band shifted slightly to 1724.36 cm⁻¹, consistent with involvement of phenolic hydroxyl and carbonyl groups in reduction of Ag+ to Ag0 [23,24]. C–H bands were retained at 2916.37/2850.79 cm⁻¹, and the C–O stretch shifted from 1053.13 to 1016.49 cm⁻¹, confirms that the flavonoid molecules effectively capped the nanoparticle surfaces [21]. These data indicate that the plant-derived flavonoids serve a dual role as both the primary reducing agents and the stabilizing ligands, ensuring the structural integrity and stability of the synthesized AgNPs.

**Figure 2.**
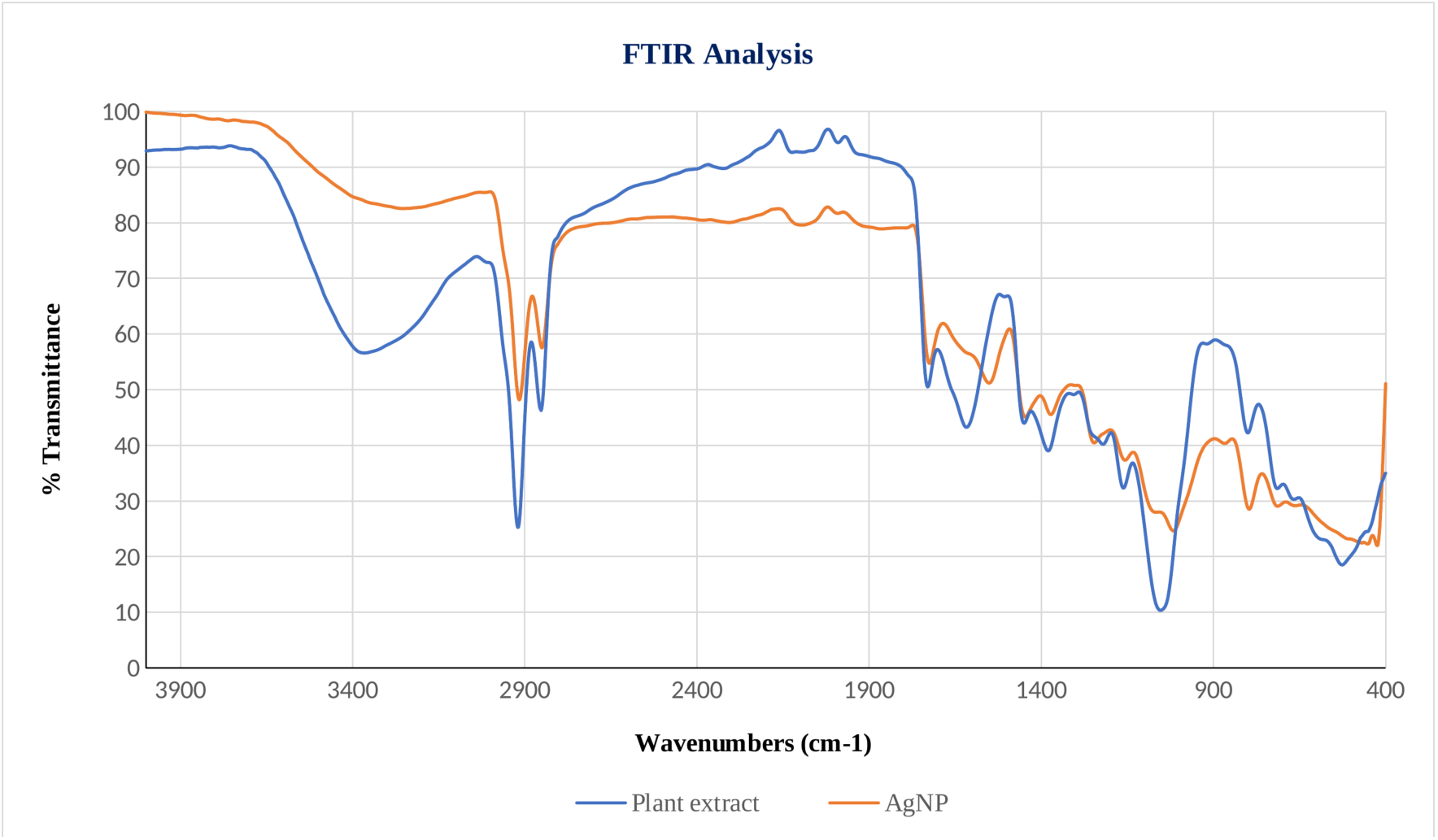
FTIR-ATR spectra of the flavonoid-enriched fraction and the green-synthesised AgNPs, showing the bathochromic shift of the O–H band and the shift of the C=O band consistent with participation of phenolic hydroxyl and carbonyl groups in reduction and surface capping.

#### 3.4.3 Zeta Potential Analysis

The surface charge and long-term colloidal stability of the green-synthesized silver nanoparticles (AgNPs) were evaluated using Laser Doppler Electrophoresis. The synthesized AgNPs exhibited a significant negative Zeta potential of -28.4 mV and a zeta deviation of 4.15 mV [25,26], with the measurement validated as "Good" quality by the Malvern analytical software (Figure 3A). The analysis was maintained at a low suspension conductivity of 0.086 mS/cm, suppressing electrical interference and Joule heating during measurement. Per the classification scheme of [27], zeta potential magnitudes in the 20–30 mV range are categorized as moderately stable. This is consistent with values reported for other green-synthesized AgNP systems in the same range, which have similarly been described as showing acceptable colloidal stability for biological applications [28].

**Figure 3.**
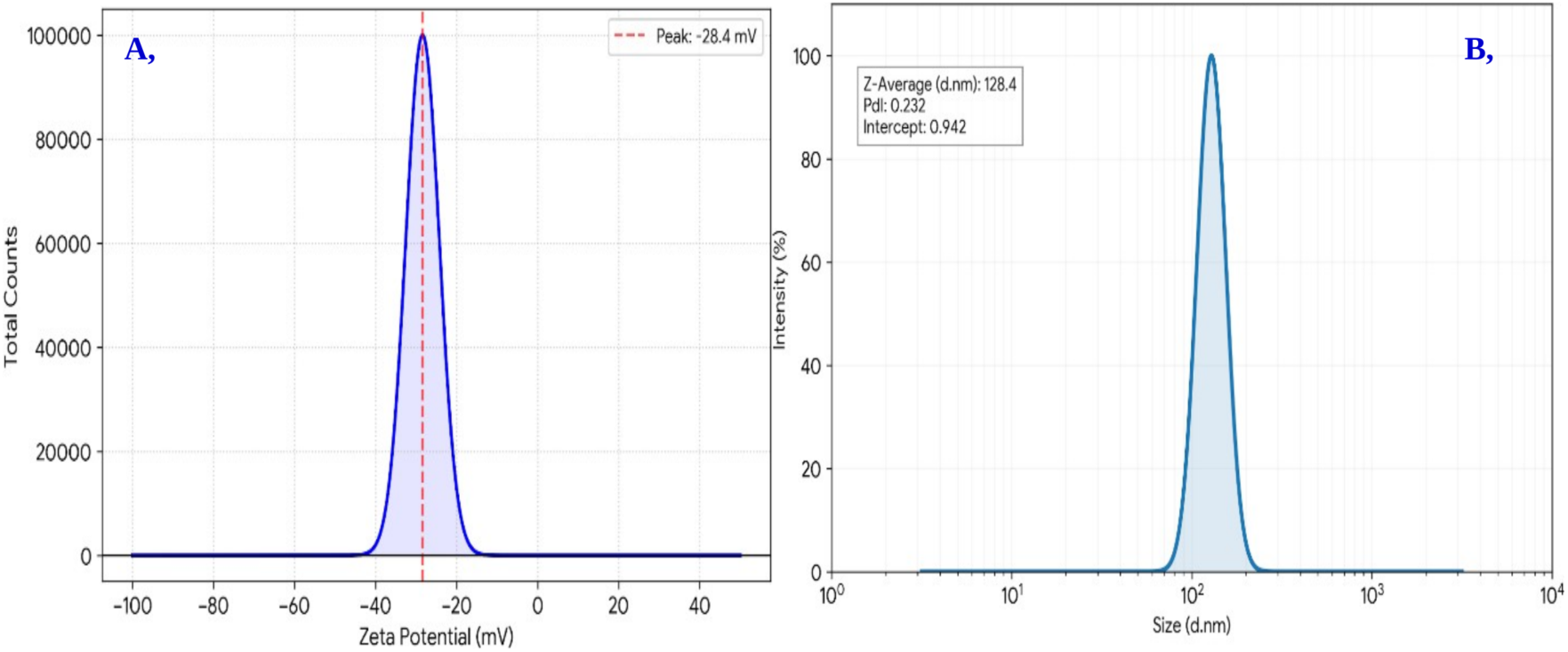
Colloidal characterisation of the green-synthesised AgNPs. (A) Zeta potential distribution measured by laser Doppler electrophoresis. (B) Hydrodynamic size distribution measured by dynamic light scattering.

#### 3.4.4 Hydrodynamic Size and Polydispersity Analysis (DLS)

The hydrodynamic diameter and the physical stability of the green-synthesized silver nanoparticles (AgNPs) were determined using Dynamic Light Scattering (DLS) [29]. The DLS analysis revealed that the AgNPs possessed a Z-average diameter of 128.4 nm and Polydispersity Index (PdI) was found to be 0.232 (Figure 3B); a PdI below 0.3 is generally considered indicative of a relatively narrow, monodisperse distribution [30]. The intercept value of 0.942 further confirms the robustness of the measurement and the absence of significant aggregation in the sample during analysis.

#### 3.4.5 Scanning Electron Microscopy (SEM) Analysis

The surface topography and structural morphology of the green-synthesized silver nanoparticles (AgNPs) were evaluated using scanning electron microscopy (Figure 4B-D). Micrographs showed predominantly spherical to sub-spherical particles distributed across an organic matrix, with discrete, well-bounded particles visible at higher magnification and localized clustering in some regions.

**Figure 4.**
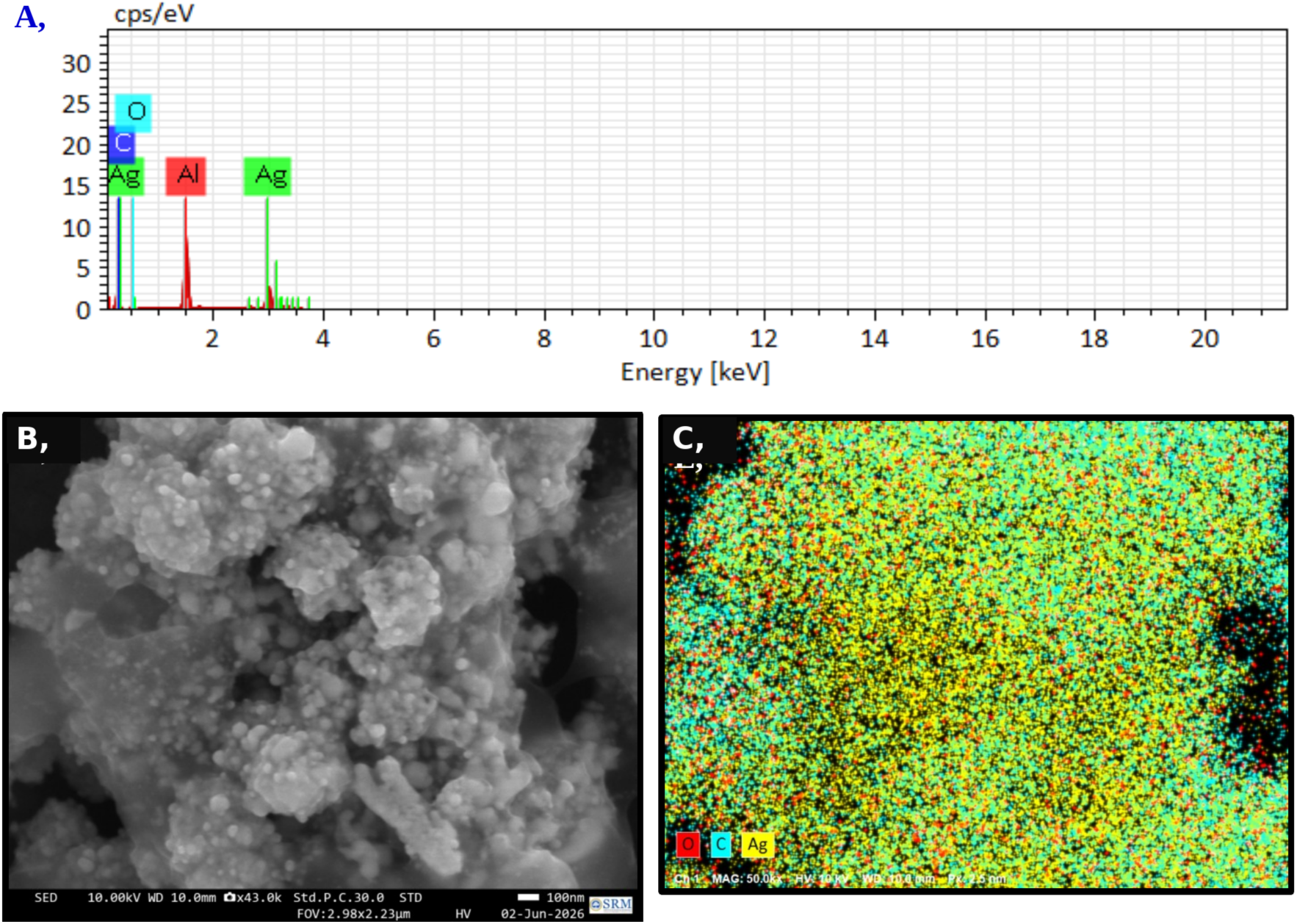
Elemental and morphological characterisation of the green-synthesised AgNPs. (A) EDS spectrum. (B) FE-SEM micrograph showing spherical to sub-spherical particles distributed across an organic matrix. (C) Elemental map showing silver co-localised with carbon and oxygen across the same regions.

#### 3.4.6 Energy-Dispersive X-ray Spectroscopy (EDS) and Elemental Mapping

Elemental composition was assessed by EDS (Figure 4A) to confirm the metallic identity of the particles observed by FE-SEM (Figure 4B; additional magnifications in Figure S3). Elemental maps (Figure 4C) showed silver co-localised with the bright topographical features, with carbon and oxygen co-localised across the same regions, consistent with a flavonoid corona surrounding the metallic cores. Quantitative analysis (Table S3) gave a normalised composition of Ag 81.26%, C 12.44% and O 6.29%. The minor aluminium peak present in the spectrum originates from the aluminium stub used for sample mounting and was therefore excluded from the final quantitative elemental analysis.

### 3.5 Acute Cytotoxicity Evaluation (WST-1 Assay)

The acute in vitro cytotoxic profiles of the flavonoid-rich fraction and its green-synthesized silver nanoparticles (AgNPs) on Vero cells were evaluated using the WST-1 colorimetric assay to determine their safe dosage range for subsequent biomedical and antiviral investigations. The flavonoid-rich fraction showed a CC₅₀ of 134.9 µg/mL (95% CI: 110.77–166.20 µg/mL) against Vero cells, while the AgNPs showed a CC50 of 1,165.74 µg/mL (95% CI: 875.21–1597.69 µg/mL), establishing a significant reduction in acute cellular toxicity characterized by an approximately 8.6-fold increase in the cellular viability threshold (Figure 5A).

**Figure 5.**
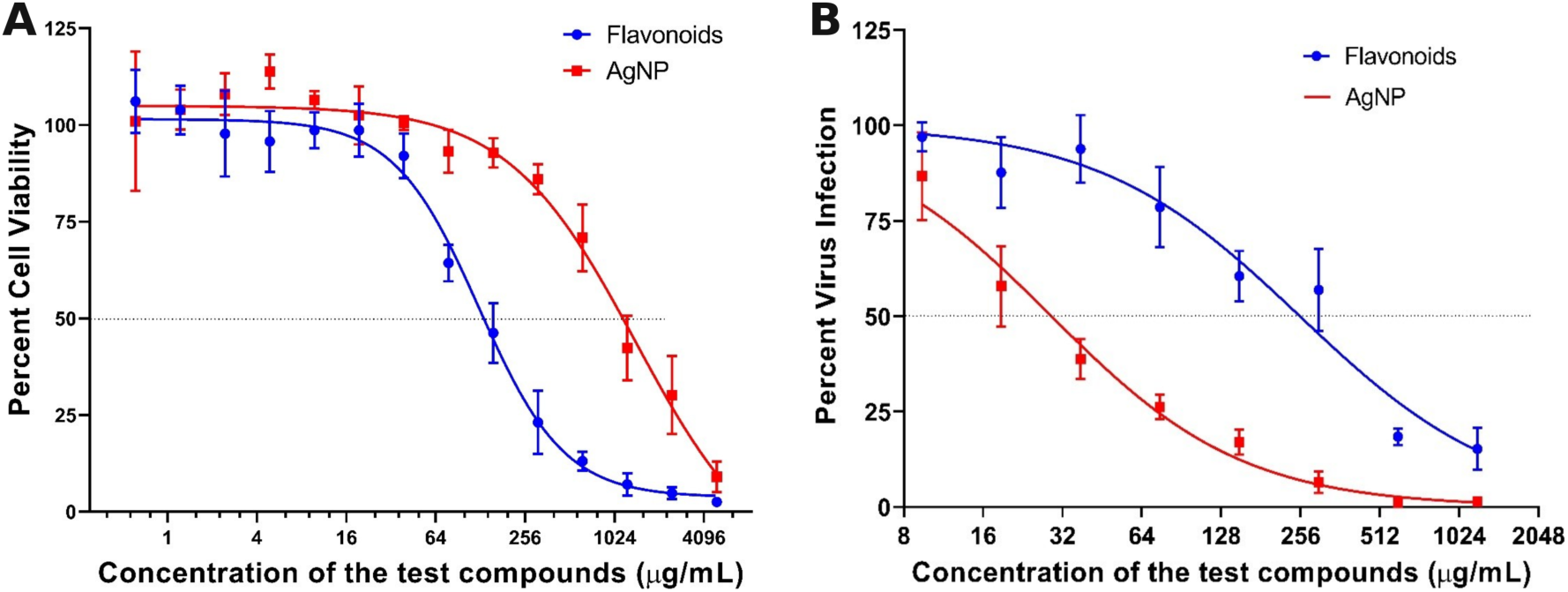
Inversion of the selectivity profile on nanoparticle formation. (A) WST-1 cytotoxicity dose-response for the flavonoid-rich fraction and the AgNPs in Vero cells. (B) Dose-dependent neutralisation of DENV-2 by the two preparations. Data are mean ± SEM of three independent replicates fitted to a four-parameter logistic model.

### 3.6. In Vitro Antiviral Efficacy and Pan-Dengue Neutralization

#### 3.6.1. Dose-Dependent Neutralization of DENV-2

The initial neutralization potential of the formulations was established using a dose-dependent pre-treatment assay against the DENV-2 serotype (Figure 5B). The green-synthesized AgNPs exhibited markedly greater antiviral activity against DENV-2 than the raw flavonoid fraction. The half-maximal effective concentration (EC₅₀) for the AgNPs was calculated at 29.05 µg/mL (95% CI: 23.40–35.90 µg/mL), whereas the flavonoid fraction required an EC₅₀ of 254.4 µg/mL (95% CI: 191.1–344.13 µg/mL). This corresponds to an approximately 8.8-fold higher antiviral potency for the AgNPs. Both dose-response models demonstrated acceptable goodness of fit (R² = 0.938 for AgNPs and 0.865 for the flavonoid fraction).

#### 3.6.2. Time-of-Addition Analysis (Mechanism of Action)

To delineate the specific stage of the viral life cycle targeted by the nanoparticles, a time-of-addition study (pre-treatment, co-treatment, and post-treatment) was conducted against DENV-2. Based on the calculated EC₅₀, a standardized inhibitory concentration of 30 µg/mL was utilized for all subsequent mechanistic assays.

The pre-treatment (neutralization) of the virus prior to cell inoculation resulted in the most significant inhibition. At 30 µg/mL, the AgNPs significantly reduced viral infectivity compared to both the untreated control (p = 0.0190) and the raw flavonoid fraction (p = 0.0012). In the co-treatment (during infection) assay, where formulations and virus were introduced to the cells simultaneously, the AgNPs successfully interfered with viral attachment and entry, again demonstrating a statistically significant reduction in infectivity relative to the control (p = 0.0363) and the raw flavonoids (p = 0.0004). In both the pre- and co-treatment models (Figure 6), the raw flavonoid fraction alone at the lower concentration 30 µg/mL failed to induce statistically significant viral inhibition compared to the control (p = 0.2557 and p = 0.6110, respectively).

**Figure 6.**
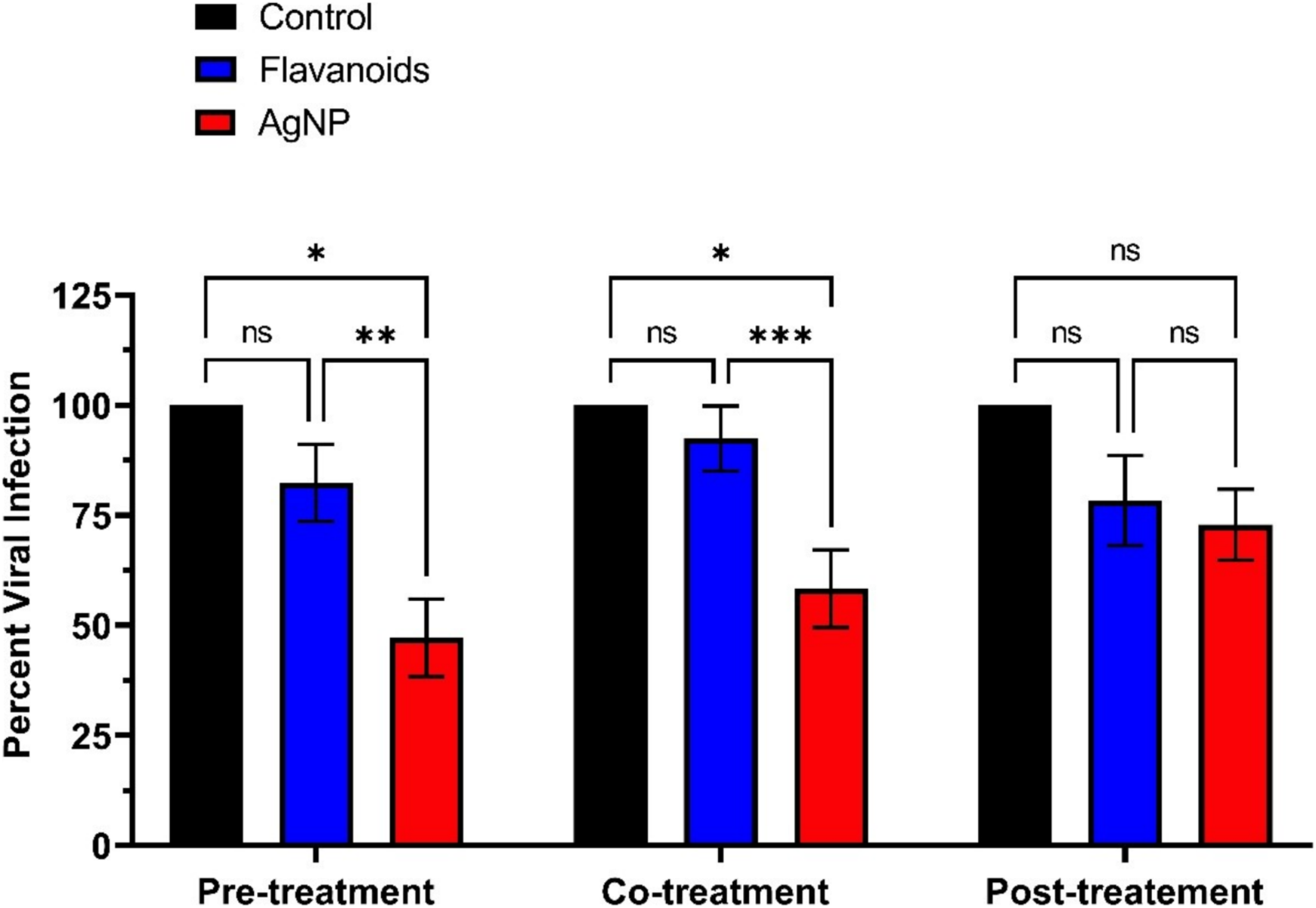
Time-of-addition analysis against DENV-2 at 30 µg/mL, comparing pre-treatment, co-treatment and post-treatment regimens. Data are mean ± SEM of three independent replicates.

Conversely, in the post-treatment assay, where formulations were introduced after a 2-hour viral adsorption period, the AgNPs reduced viral infection by approximately 27%; however, this trend did not reach statistical significance relative to the untreated control (p = 0.0850). Similarly, the raw flavonoids failed to induce a significant reduction (p = 0.2321). These results indicate that the AgNPs act primarily as potent therapeutic entry inhibitors and systemic neutralizing agents, intercepting the virus prior to or during host-cell attachment, rather than targeting post-entry intracellular replication.

#### 3.6.3. Broad-Spectrum (Pan-Dengue) Efficacy

To investigate the broad-spectrum (pan-dengue) potential of the formulations, efficacy was further evaluated against the remaining three serotypes (DENV-1, DENV-3, and DENV-4) along with DENV-2. Utilizing the established 30 µg/mL concentration as a standardized baseline, cross-serotype susceptibility was compared.

As shown Figure 7, the raw flavonoid fraction at 30 µg/mL exhibited negligible activity against DENV-1, -2 and -4; a modest but statistically significant reduction was observed for DENV-3 (∼21%), still substantially inferior to the AgNPs. In stark contrast, the AgNPs at the identical concentration (30 µg/mL) demonstrated robust, broad-spectrum neutralization. While the AgNPs reduced DENV-2 infectivity by approximately 50% (consistent with its established EC₅₀), they exhibited even greater neutralizing potency against other serotypes. DENV-1 proved to be the most susceptible to the AgNPs, with viral infectivity dropping to approximately 21% (representing an ∼79% viral inhibition). Similarly, DENV-4 and DENV-3 were profoundly neutralized, with residual viral infectivities reduced to approximately 24% and 39%, respectively. Collectively, these results confirm that the green-synthesized AgNPs possess potent pan-dengue antiviral properties, outperforming the raw extract across all four viral serotypes.

**Figure 7.**
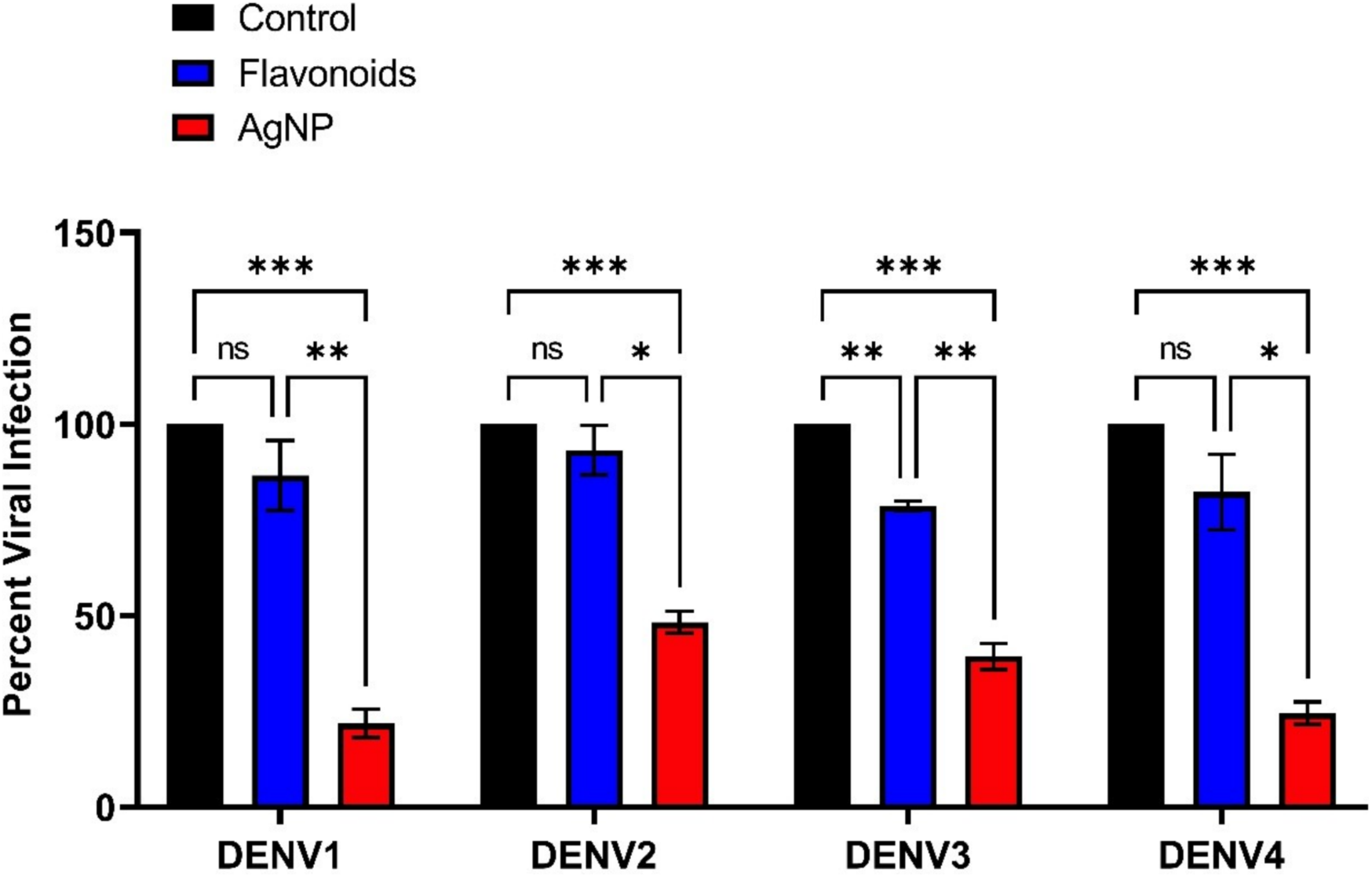
Cross-serotype screen at 30 µg/mL against DENV-1, DENV-2, DENV-3 and DENV-4, expressed as residual viral infectivity relative to the untreated control. Data are mean ± SEM of three independent replicates.

### 3.7 Selectivity index and Safety Margin Evaluation

The clinical safety and applicability of both the Carica papaya flavonoid fraction and its synthesized AgNPs were quantified using the Selectivity index (SI), where a value greater than 10 establishes a safe and promising window for therapeutic drug development (De Clercq, 2004).

The flavonoid-rich fraction gave a CC₅₀ of 134.9 µg/mL against an EC₅₀ of 254.4 µg/mL, yielding a Selectivity Index of 0.53. The green-synthesised AgNPs gave a CC₅₀ of 1,165.74 µg/mL against an EC₅₀ of 29.05 µg/mL, yielding a Selectivity Index of 40.12. Formulation therefore shifted the selectivity profile approximately 75-fold, from a value below unity, at which no therapeutic window exists, to one above the threshold conventionally applied in antiviral screening.

## 4. DISCUSSION

### 4.1 Phytochemical Basis for Bioactivity

The chloroform fraction’s high flavonoid content (55.79 ± 1.94 mg QE/g) is consistent with the well-documented antioxidant, membrane-interacting, and virucidal properties reported for flavonoids against enveloped RNA viruses [31] and supports its selection as the reducing/capping matrix for AgNP synthesis.

### 4.2 Mechanistic Basis of Green Synthesis and Stabilization

The spectroscopic and surface-charge data collectively support a green-synthesis mechanism in which C. papaya flavonoids act dually as reducing and capping agents. The transition from the extract’s UV-Vis absorption band (287 nm) to the AgNP SPR peak (417 nm) reflects electron donation from flavonoid hydroxyl/carbonyl groups during Ag⁺ reduction to Ag⁰. The FTIR shifts observed at the O–H and C=O bands corroborate this, indicating that the same functional groups responsible for reduction subsequently form the capping layer that confers colloidal stability, consistent with the DLS/zeta-potential profile (moderately stable, as per section 3.4.3) and the SEM/EDS evidence of an organic corona surrounding the silver cores [32,33].

### 4.3 Cytotoxicity Reduction Following Nanoparticle Conversion

The approximately 8.6-fold increase in CC₅₀ upon conversion of the raw fraction into AgNPs (134.9 → 1,165.74 µg/mL) is consistent with a mechanism in which the biogenic capping corona moderates direct cellular exposure to free phytochemicals and limits uncontrolled intracellular silver-ion leaching, thereby reducing acute oxidative stress during the 24-hour exposure period [34]. This safety profile compares favorably to cytotoxicity profiles reported elsewhere in the green-synthesis literature [35].

### 4.4 Antiviral Mechanism of Action, Broad-Spectrum Efficacy, and Clinical Relevance

The substantially greater potency of the AgNPs relative to the raw fraction (EC₅₀ = 29.05 vs. 254.4 µg/mL, an ∼8.8-fold enhancement) suggests that nanoparticle conversion actively potentiates the flavonoids’ intrinsic antiviral activity. Flavonoids such as quercetin and kaempferol are known to interfere with DENV, but their therapeutic use is often constrained by poor aqueous-phase stability and limited bioavailability [36]. This limitation is circumvented here by anchoring the bioactive molecules to a stable metallic carrier. The silver core effectively acts as a multivalent scaffold, increasing the local concentration and surface density of the flavonoid molecules and presenting them in an organized configuration that facilitates high-avidity interactions with the viral surface [37].

The specific stage of the viral life cycle targeted by this multivalent platform was definitively elucidated through the Time-of-Addition analysis. The AgNPs demonstrated their most profound and statistically significant viral inhibition during the pre-treatment and co-treatment (during infection) stages. When introduced post-infection, the formulation retained partial efficacy (reducing viral infectivity by approximately 27% at the low 30 µg/mL dose); however, the data clearly indicates that its primary, most potent action occurs prior to or during host-cell attachment. Critically, this extracellular entry-inhibition represents a major clinical advantage. Pathogenesis in severe dengue is driven by high levels of acute viremia, which triggers the cytokine cascades responsible for plasma leakage. By neutralizing the virus in the extracellular space, this AgNPs can function mechanistically similar to clinical monoclonal antibodies (mAbs), a circulating viral trap that clears viremia from the bloodstream.

However, while therapeutic monoclonal antibodies (mAbs) are highly effective in other viral diseases, their clinical translation for Dengue is severely constrained by high production costs, cold-chain storage requirements, and the critical risk of Antibody-Dependent Enhancement (ADE) if cross-serotype neutralization is incomplete or sub-optimal [4,38]. The biogenic AgNP platform overcomes these fundamental demerits. Because the AgNPs rely on broad-spectrum physical mechanisms and surface disruption rather than single-epitope recognition, they theoretically mitigate the Fc-receptor-mediated risks of ADE, while their inherent thermodynamic stability and low-cost green synthesis offer a highly scalable alternative for endemic regions [39].

Furthermore, a viable clinical therapeutic for Dengue must address the co-circulation of its multiple serotypes. The raw flavonoid fraction at 30 µg/mL lacked the potency to induce meaningful neutralization across the viral family. In contrast, the formulated AgNPs at the identical concentration demonstrated potent, broad-spectrum (pan-dengue) neutralization against DENV-1, DENV-2, DENV-3, and DENV-4. Two non-exclusive mechanisms likely account for this cross-serotype efficacy: (i) steric hindrance, where the nanoparticles physically occlude the E-protein from interacting with host cellular receptors, and (ii) structural disruption induced by the highly reactive flavonoid corona [40]. Collectively, these mechanistic and broad-spectrum data elevate the green-synthesized AgNPs from a simple botanical extract to a highly potent, systemic nanotherapeutic candidate for acute dengue intervention.

### 4.5 Selectivity index and Safety Margin

The conversion of the raw flavonoid fraction into AgNPs resulted in a profound expansion of the safety margin, with the Selectivity index (SI) improving from a Cytotoxically limiting 0.53 (CC₅₀ 134.9 / EC₅₀ 254.4) to a safe and highly viable 40.12 (CC₅₀ 1165.74 / EC₅₀ 29.05). This ∼75-fold enhancement easily clears the established SI ≥ 10 threshold conventionally required for viable antiviral drug candidates [41]. This improvement appears attributable to two related effects: first, the capping layer’s role in moderating cytotoxicity (as per section 4.3) raises the CC₅₀ denominator; second, the organized surface presentation of flavonoids on the AgNP scaffold appears to lower the concentration needed for antiviral effect, reducing the EC₅₀ numerator [12]. It is worth noting that this SI reflects a 24-hour exposure model; some other phytosynthesized metal nanoparticle systems report selectivity indices in the range of approximately 5–20 under different exposure conditions [42].

### 4.6 Limitations

This study has some limitations that should inform interpretation of the findings. Both the cytotoxicity and Selectivity Index values reflect a 24-hour exposure paradigm; chronic or repeated-exposure toxicity profiles may differ substantially and would need separate evaluation before any translational claims. Additionally, a canonical antiviral comparator was not applicable here given the direct virucidal/entry-inhibition mechanism observed, and an uncapped ionic silver control was not pursued as it is not chemically equivalent to the phytochemical-capped nanoparticle under study. Finally, all findings are based on in vitro assays; in vivo efficacy, pharmacokinetics, and biodistribution of the AgNP formulation remain to be established.

## 5. CONCLUSION

This study demonstrates that the green synthesis of silver nanoparticles using a Carica papaya flavonoid fraction effectively rescues a botanically active but otherwise cytotoxically limiting phyto-extract, transforming it into a potent and safer, pan-dengue nanotherapeutic. By utilizing the flavonoids as dual (reducing and capping) agents, the resulting AgNP formulation exhibited an ∼8.6-fold reduction in cellular toxicity alongside an ∼8.8-fold enhancement in antiviral potency. This dual action dramatically expanded the Selectivity Index from an unviable 0.53 to a safe margin of 40.12. Mechanistically, these nanoparticles function as potent therapeutic entry inhibitors. By neutralizing the virus prior to and during host-cell attachment, they can act as circulating "viral traps" capable of addressing acute viremia across all four dengue serotypes (DENV 1–4) comparable to the circulating neutralizing antibodies. Because of their cross-serotype activity, these biogenic AgNPs may overcome the serotype-specific limitations and potential ADE risks associated with conventional antibody-based therapeutics and vaccines. Their physical mode of viral neutralization further highlights their potential as a scalable candidate for acute dengue intervention. Future in vivo pharmacokinetic and efficacy investigations are highly warranted to advance this botanical nanotherapeutic toward clinical application.

## Ethics and biosafety approval

The experimental protocols and study design were formally reviewed and approved by the Institutional Ethics Committee (Registration No. ECR/8853/INST/TN/2013/RR-19). Furthermore, all procedures involving live viral strains and in vitro cell culture were conducted in strict accordance with biosafety guidelines and received formal approval from the Institutional Biosafety Committee (IBSC Ref. No. TFR:NCBS:46IBSC/LS/6/N) of NCBS.

## Author contributions

Madan Kumar D: Conceptualization, Methodology, Investigation, Data curation, Formal analysis, Visualization, Writing – original draft. R. Raj Bharath: Supervision, Project administration, Writing – review and editing. Gowtham Palanisamy: Resources. G. Devanand Venkatasubbu: Resources. Tanay Bhatt: Methodology, Resources, Supervision, Writing – review and editing.

## Data availability

The data supporting the findings of this study are available from the corresponding author upon reasonable request.

## Supplementary information

Supplementary Figures S1 to S3 and Supplementary Tables S1 to S3 are provided as a separate file.

## Acknowledgements

The authors would like to express their sincere gratitude to Prof. L.S. Shashidhara, Centre Director, National Centre for Biological Sciences (NCBS), Tata Institute of Fundamental Research, Bangalore, for generously providing the specialized biosafety laboratory infrastructure at NCBS necessary to conduct the cellular and viral assays. The authors gratefully acknowledge the Nanotechnology Research Centre (NRC) and the SRM Central Instrumentation Facility (SCIF) at SRMIST for providing the state-of-the-art diagnostic platforms utilized for the structural, morphological, and optical characterization of the green-synthesized AgNPs. Sincere gratitude is extended to Simbioens Laboratory for providing vital experimental research facilities.

## Declaration of generative AI and AI-assisted technologies in the manuscript preparation process

During the preparation of this work the authors used Claude (Anthropic) in order to assist with language editing and manuscript structuring. After using this tool, the authors reviewed and edited the content as needed and take full responsibility for the content of this manuscript.

## References

[1] S. Bhatt, P.W. Gething, O.J. Brady, J.P. Messina, A.W. Farlow, C.L. Moyes, et al., The global distribution and burden of dengue, Nature 496, 504–507 (2013). 10.1038/nature12060

2. WHO, DENGUE GUIDELINES FOR DIAGNOSIS, TREATMENT, PREVENTION AND CONTROL TREATMENT, PREVENTION AND CONTROL TREATMENT, PREVENTION AND CONTROL, www.who.int/tdr (2009).

[3] J.D. Stanaway, D.S. Shepard, E.A. Undurraga, Y.A. Halasa, L.E. Coffeng, O.J. Brady, et al., The global burden of dengue: an analysis from the Global Burden of Disease Study 2013, The Lancet Infectious Diseases 16, 712–723 (2016). 10.1016/S1473-3099(16)00026-8

[4] A. Sarker, N. Dhama, R.D Gupta, Dengue virus neutralizing antibody: a review of targets, cross-reactivity, and antibody-dependent enhancement, Frontiers in Immunology, 14 (2023). 10.3389/fimmu.2023.1200195

5. World Health Organization, Dengue, (2025). https://www.who.int/news-room/fact-sheets/detail/dengue-and-severe-dengue

[6] S. Sridhar, A. Luedtke, E. Langevin, M. Zhu, M. Bonaparte, T. Machabert, et al., Effect of Dengue Serostatus on Dengue Vaccine Safety and Efficacy, New England Journal of Medicine 379, 327–340 (2018). 10.1056/NEJMoa1800820

[7] K. Zandi, B.-T. Teoh, S.-S. Sam, P.-F. Wong, M.R. Mustafa, S AbuBakar, Antiviral activity of four types of bioflavonoid against dengue virus type-2, Virology Journal 8, 560 (2011). 10.1186/1743-422X-8-560

[8] B.E. Dewi, H. Desti, E. Ratningpoeti, M. Sudiro, Fithriyah, M Angelina, Effectivity of quercetin as antiviral to dengue virus-2 strain New Guinea C in Huh 7-it 1 cell line, IOP Conference Series: Earth and Environmental Science 462, 012033 (2020). 10.1088/1755-1315/462/1/012033

[9] T. Nguyen, M.-O. Parat, M. Hodson, J. Pan, P. Shaw, A Hewavitharana, Chemical Characterization and in Vitro Cytotoxicity on Squamous Cell Carcinoma Cells of Carica Papaya Leaf Extracts, Toxins 8, 7 (2015). 10.3390/toxins8010007

[10] A. Luceri, R. Francese, D. Lembo, M. Ferraris, C Balagna, Silver Nanoparticles: Review of Antiviral Properties, Mechanism of Action and Applications, Microorganisms 11, 629 (2023). 10.3390/microorganisms11030629

[11] M. Moond, S. Singh, S. Sangwan, R. Devi, R Beniwal, Green Synthesis and Applications of Silver Nanoparticles: A Systematic Review, AATCC Journal of Research 9, 272–285 (2022). 10.1177/24723444221119847

[12] A.W. Bere, O. Mulati, J. Kimotho, F Ng’ong’a, Carica papaya Leaf Extract Silver Synthesized Nanoparticles Inhibit Dengue Type 2 Viral Replication In Vitro, Pharmaceuticals 14, 718 (2021). 10.3390/ph14080718

13. W.C Evans, Trease and Evans’ pharmacognosy, Elsevier Health Sciences (2009).

[14] J.B Harborne, Phytochemical Methods: A Guide to Modern Techniques of Plant Analysis, (3rd ed.). Chapman & Hall (1998).

[15] M. Iqbal, A.A Jariah, Phytochemical screening of secondary metabolite compounds in tammate leaf extract (Lannea coromandelica (HOUTT,) MERR.) from Pangkep Regency using various extraction methods. Journal of Current Health Sciences 5, 59–66 (2025). 10.47679/jchs.2025107

[16] C.-C. Chang, M.-H. Yang, H.-M. Wen, J.-C Chern, Estimation of total flavonoid content in propolis by two complementary colometric methods, Journal of Food and Drug Analysis 10 (2020). 10.38212/2224-6614.2748

[17] R.R. Banala, V.B. Nagati, P.R Karnati, Green synthesis and characterization of Carica papaya leaf extract coated silver nanoparticles through X-ray diffraction, electron microscopy and evaluation of bactericidal properties, Saudi Journal of Biological Sciences 22, 637–644 (2015). 10.1016/j.sjbs.2015.01.007

[18] A.H. Labulo, O.A. David, A.D Terna, Green synthesis and characterization of silver nanoparticles using Morinda lucida leaf extract and evaluation of its antioxidant and antimicrobial activity, Chemical Papers 76, 7313–7325 (2022). 10.1007/s11696-022-02392-w

[19] G. Añez, E. Volkova, Z. Jiang, D.A.R. Heisey, C. Chancey, R.C.G. Fares, et al., Collaborative study to establish World Health Organization international reference reagents for dengue virus Types 1 to 4 RNA for use in nucleic acid testing, Transfusion 57, 1977–1987 (2017). 10.1111/trf.14130

[20] G.A. Santiago, E. Vergne, Y. Quiles, J. Cosme, J. Vazquez, J.F. Medina, et al., Analytical and Clinical Performance of the CDC Real Time RT-PCR Assay for Detection and Typing of Dengue Virus, PLoS Neglected Tropical Diseases 7, e2311 (2013). 10.1371/journal.pntd.0002311

[21] A.K. Alzubaidi, W.J. Al-Kaabi, A. Al Ali, S. Albukhaty, H. Al-Karagoly, G.M. Sulaiman, et al., Green Synthesis and Characterization of Silver Nanoparticles Using Flaxseed Extract and Evaluation of Their Antibacterial and Antioxidant Activities, Applied Sciences 13, 2182 (2023). 10.3390/app13042182

[22] M.R. Siddiqui, M. Khan, Khan, Adil, Tahir, W. Tremel, et al., Green synthesis of silver nanoparticles mediated by Pulicaria glutinosa extract, International Journal of Nanomedicine, 1507 (2013). 10.2147/IJN.S43309

[23] A. Purohit, R. Sharma, R. Shiv Ramakrishnan, S. Sharma, A. Kumar, D. Jain, et al., Biogenic Synthesis of Silver Nanoparticles (AgNPs) Using Aqueous Leaf Extract of Buchanania lanzan Spreng and Evaluation of Their Antifungal Activity against Phytopathogenic Fungi, Bioinorganic Chemistry and Applications 2022 (2022). 10.1155/2022/6825150

[24] R. Das, P. Kumar, A.K. Singh, S. Agrawal, S. Albukhaty, I. Bhattacharya, et al., Green synthesis of silver nanoparticles using Trema Orientalis (L,) extract and evaluation of their antibacterial activity. Green Chemistry Letters and Reviews 18 (2025). 10.1080/17518253.2024.2444679

[25] A.T.M. Saeb, A.S. Alshammari, H. Al-Brahim, K.A Al-Rubeaan, Production of Silver Nanoparticles with Strong and Stable Antimicrobial Activity against Highly Pathogenic and Multidrug Resistant Bacteria, The Scientific World Journal 2014, 1–9 (2014). 10.1155/2014/704708

[26] D. Satria, N.A. Siregar, S.B. Waruwu, E.D.L. Putra, S.M. Sinaga, A.W. Septama, et al., Eco-friendly one-pot synthesis of silver nanoparticles via microwave irradiation using Artocarpus camansi extract: Characterization, antioxidant, and antibacterial applications, Journal of King Saud University – Science 38, 10112025 (2026). 10.25259/JKSUS_1011_2025

[27] S Bhattacharjee, DLS and zeta potential – What they are and what they are not? Journal of Controlled Release, 235, 337–351, (2016). 10.1016/j.jconrel.2016.06.017

[28] N.P. Lemeitaron, M.E. Shigwenya, L.N. Munuhe, S.D. Makhanu, K.P Kinyanjui, Green synthesis of silver nanoparticles via aqueous extracts of Prunus africana and their antimicrobial activities, South African Journal of Chemical Engineering 57, 100883 (2026). 10.1016/j.sajce.2026.100883

[29] A.K.O. Lima, L.M. dos S. Souza, G.F. Reis, A.G.T. Junior, V.H.S. Araújo, L. C. dos Santos, et al., Synthesis of Silver Nanoparticles Using Extracts from Different Parts of the Paullinia cupana Kunth Plant: Characterization and In Vitro Antimicrobial Activity, Pharmaceuticals 17, 869 (2024). 10.3390/ph17070869

[30] M. Danaei, M. Dehghankhold, S. Ataei, F. Hasanzadeh Davarani, R. Javanmard, A. Dokhani, et al., Impact of Particle Size and Polydispersity Index on the Clinical Applications of Lipidic Nanocarrier Systems, Pharmaceutics 10, 57 (2018). 10.3390/pharmaceutics10020057

[31] I. Sánchez, F. Gómez-Garibay, J. Taboada, B.H Ruiz, Antiviral effect of flavonoids on the Dengue virus, Phytotherapy Research 14, 89–92 (2000). 10.1002/(SICI)1099-1573(200003)14:2<89::AID-PTR569>3.0.CO;2-C

[32] Y. Han, L. Zhu, X. Zhang, K. Wang, D. Sha, Q. Chen, et al., Green synthesis, characterization, and anticancer activity of silver nanoparticles via Pulsatilla koreana root extract, Frontiers in Nutrition, 13 (2026). 10.3389/fnut.2026.1746280

[33] E.O Mikhailova, Green Silver Nanoparticles: An Antibacterial Mechanism, Antibiotics 14, 5 (2024). 10.3390/antibiotics14010005

[34] J. Jeevanandam, S. Krishnan, Y.S. Hii, S. Pan, Y.S. Chan, C. Acquah, et al., Synthesis approach-dependent antiviral properties of silver nanoparticles and nanocomposites, Journal of Nanostructure in Chemistry 12, 809–831 (2022). 10.1007/s40097-021-00465-y

[35] S.F. hamieda, M Saied, In Vitro Evaluation of Cytotoxicity and Antimicrobial Activity of Green Synthesized Silver Nanoparticles Based on Their Particle Size and Stability, Egyptian Journal of Chemistry 68, 79–89 (2025). 10.21608/ejchem.2024.328372.10631

[36] S. Thilakarathna, H Rupasinghe, Flavonoid Bioavailability and Attempts for Bioavailability Enhancement, Nutrients 5, 3367–3387 (2013). 10.3390/nu5093367

[37] F.S. Hussain, N.Q. Abro, N. Ahmed, S.Q. Memon, N Memon, Nano-antivirals: A comprehensive review, Frontiers in Nanotechnology, 4 (2022). 10.3389/fnano.2022.1064615

[38] D. G. vander Sterren, I., vanLeeuwen, L.P.M. Aynekulu Mersha, T. Langerak, M.S. Hakim, B., van Lelyveld, S.F.L. Martina, E.C.M van Gorp, The role of antibody-dependent enhancement in dengue vaccination, Tropical Diseases, Travel Medicine and Vaccines 10, 22 (2024). 10.1186/s40794-024-00231-2

[39] Saad Ahmad, Waqar Ahmad, Danyal Ahmad, GREEN SYNTHESIS OF SILVER NANOPARTICLES AND THEIR APPLICATIONS, Kashf Journal of Multidisciplinary Research 2, 57–69 (2025). 10.71146/kjmr600

[40] A. Ahsan, G.F. Gao, W.-X Tian, Phytogenic Silver Nanoparticles Derived from Ricinus communis and Aloe barbadensis: Synthesis, Characterization, and Evaluation of Biomedical Potential, Bioengineering 12, 1273 (2025). 10.3390/bioengineering12111273

[41] E De Clercq, Antiviral drugs in current clinical use, Journal of Clinical Virology 30, 115– 133 (2004). 10.1016/j.jcv.2004.02.009

[42] M.Z. Sayed-Ahmed, M.A. Rizk, S.A.A. Hagras, M. Alfarhan, A.A. Alshamrani, A.H. Albariqi, et al., Antiviral Potential Efficacy of Green-Synthesized Silver and Titanium Dioxide Nanoparticles Against Rotavirus, Cytomegalovirus, and Human Papillomavirus, Pharmaceuticals 19, 556 (2026). 10.3390/ph19040556

