## Supplementary information for "Biogenic flavonoid capping converts a cytotoxic Carica papaya fraction into a selective, cross-serotype dengue entry inhibitor"

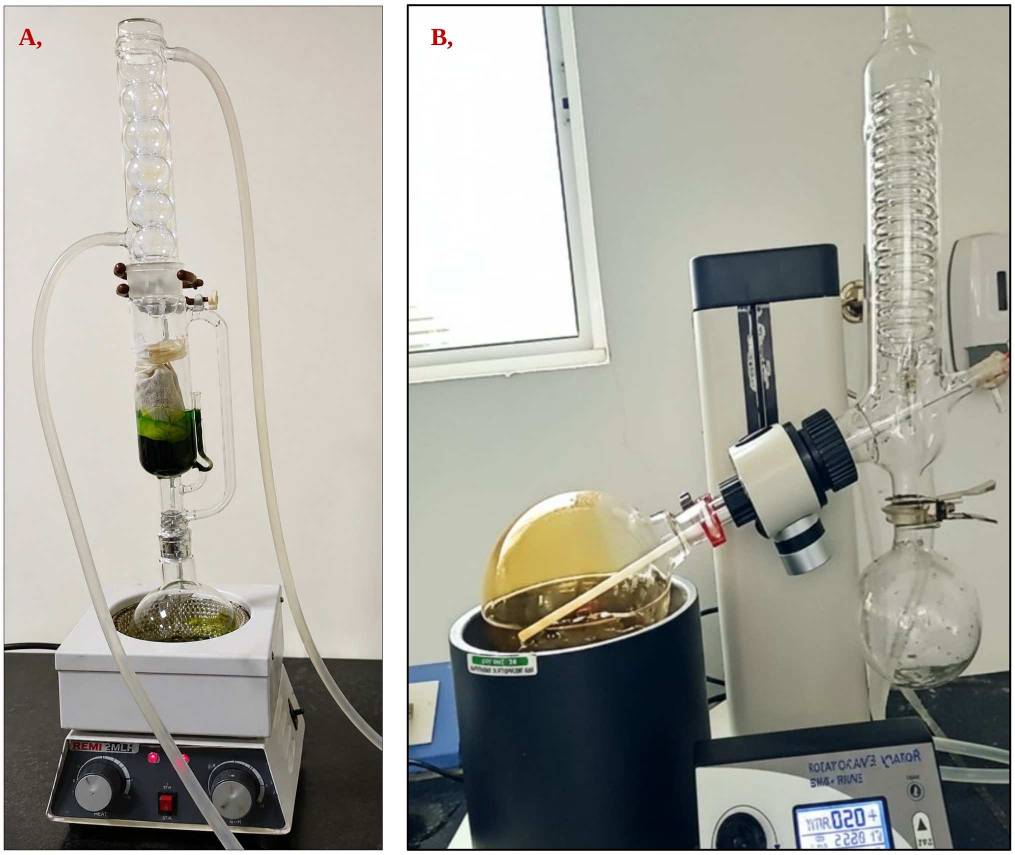


**Figure S1.** Preparation of the Carica papaya leaf extract. (A) Soxhlet extraction of shade-dried leaf powder in 70% ethanol over 52 hours. (B) Concentration of the resulting crude extract using a rotary vacuum evaporator at 40 °C.

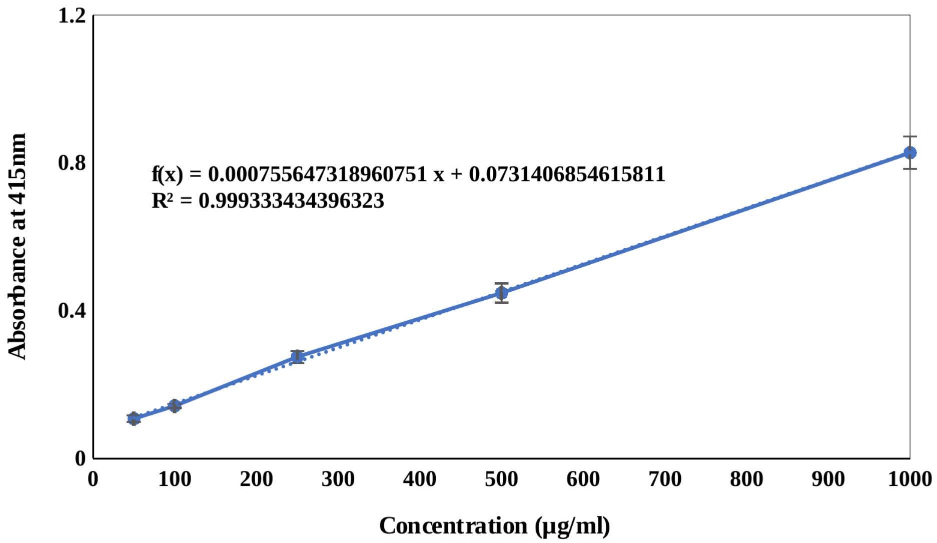


**Figure S2.** Quercetin standard calibration curve used for quantification of total flavonoid content by the aluminium chloride colorimetric method, recorded at 415 nm (y = 0.0008x + 0.0731, $R^2$ = 0.99).

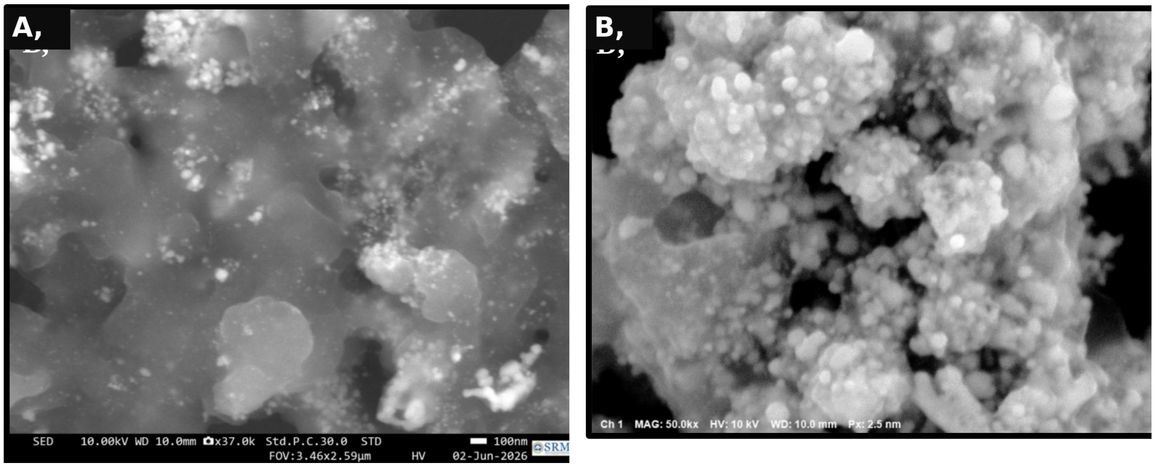


**Figure S3.** Additional FE-SEM micrographs of the green-synthesised silver nanoparticles at (A) ×37,000 and (B) ×50,000 magnification, showing spherical to sub-spherical particles with localised clustering across the organic matrix.

**Table S1.** Forward and reverse primers and hydrolysis probes for dengue virus serotypes 1 to 4, adapted from a previously validated diagnostic protocol.

| **Oligonucleotide** | **Sequence (5′ → 3′)** |
|---|---|
| DEN1_F | CAAAAGGAAGTCGYGCAATA |
| DEN1_R | CTGAGTGAATTCTCTCTGCTRAAC |
| DEN1_P | [FAM]CATGTGGYTGGGAGCRCGC[BHQ1] |
| DEN2_F | CAGGCTATGGCACYGTCACGAT |
| DEN2_R | CCATYTGCAGCARCACCATCTC |
| DEN2_P | [FAM]CTCYCCRAGAACGGGCCTCGACTTCAA[BHQ1] |
| DEN3_F | GGACTRGACACACGCACCCA |
| DEN3_R | CATGTCTCTACCTTCTCGACTTGYCT |
| DEN3_P | [FAM]ACCTGGATGTCGGCTGAAGGAGCTTG[BHQ1] |
| DEN4_F | TTGTCCTAATGATGCTRGTCG |
| DEN4_R | TCCACCYGAGACTCCTTCCA |
| DEN4_P | [FAM]TYCCTACYCCTACGCATCGCATTCCG[BHQ1] |

**Table S2.** Qualitative phytochemical screening of the 70% ethanolic crude extract of C. papaya leaves. ++ strongly present, + present, − not detected.

| **Phytochemical class** | **Result** |
|---|---|
| Flavonoids | ++ |
| Saponins | − |
| Alkaloids | − |
| Tannins | + |
| Terpenes | − |
| Phenols | + |
| Glycosides | − |
| Steroids | + |

**Table S3.** Energy-dispersive X-ray spectroscopy elemental composition of the green-synthesised silver nanoparticles. The minor aluminium signal arising from the sample-mounting stub was excluded from quantification.

| **Element** | **Z** | **Net counts** | **Mass conc. (%)** | **Norm. mass (%)** | **Norm. atomic (%)** | **3σ (mass%)** |
|---|---|---|---|---|---|---|
| Ag | 47 | 16818 | 64.29 | 81.26 | 34.51 | 9.16 |
| C | 6 | 5262 | 9.84 | 12.44 | 47.46 | 0.59 |
| O | 8 | 906 | 4.98 | 6.29 | 18.02 | 0.65 |
| Sum | | | 79.12 | 100.00 | 100.00 | |
